# MetaPepticon: automated prediction of anticancer peptides from microbial genomes and metagenomes

**DOI:** 10.1101/2025.10.28.685052

**Authors:** Ahmet Arıhan Erözden, Nalan Tavşanlı, Gamze Demirel, Nazmiye Özlem Şanlı, Mahmut Çalışkan, Muzaffer Arıkan

## Abstract

Anticancer peptides (ACPs) are increasingly recognized as promising therapeutic candidates due to their ability to selectively target cancer cells. However, the systematic discovery of novel ACPs, particularly from high-throughput sequencing datasets, remains hindered by technical and methodological limitations. Current prediction frameworks require pre-extracted peptide sequences, involve manual preprocessing, and yield variable results, which restricts their applicability for large-scale, data-driven discovery.

To address these limitations, we developed MetaPepticon, a modular, end-to-end pipeline for the discovery of candidate ACPs from diverse sequencing inputs, including raw genomic, metagenomic, transcriptomic, and metatranscriptomic reads, as well as assembled contigs and peptide sequences. MetaPepticon automates quality control, filtering, assembly, small open reading frame prediction, ACP classification using multiple predictive algorithms, and *in silico* toxicity filtering. By employing a consensus-based strategy and supporting heterogeneous data types, MetaPepticon facilitates scalable, reproducible, and high-confidence identification of candidate ACPs.

Applied to 41,171 microbial genomes and 4,072,884 peptides, MetaPepticon identified 79,587 novel candidate ACPs, including 13,149 high-confidence, non-toxic peptides. By providing a standardized, automated framework for large-scale ACP discovery across various input types, MetaPepticon facilitates therapeutic peptide exploration and is freely available at: https://github.com/arikanlab/MetaPepticon

## Background

Cancer remains the second leading cause of death worldwide, following cardiovascular diseases, and is responsible for one in every eight deaths [1]. By 2070, breast and colorectal cancer cases are expected to reach 9.1 million, representing a 131% increase from 2018, driven by demographic changes and rising incidence rates [2]. The limitations and adverse effects associated with conventional cancer treatments have driven the search for novel therapeutic alternatives [3].

Anticancer peptides (ACPs) are short peptide sequences, typically ranging from 10 to 50 amino acids, that selectively target tumor cells and induce cell death through diverse mechanisms [4,5]. Compared to traditional chemotherapy drugs, ACPs offer significant advantages, including enhanced membrane permeability, a broader range of molecular targets, and reduced adverse effects [6]. These features make ACPs promising alternatives for cancer treatment, increasing interest in their discovery and validation for therapeutic development. Consequently, computational tools for *in silico* prediction and characterization of ACPs have been increasingly developed over the past decade, facilitating initial screening and discovery of novel therapeutic peptides [6,7].

Microbial communities are rich sources of functional peptides, including ACPs [8–10], and advances in sequencing technologies have enabled large-scale exploration of their diversity, producing vast meta-omics datasets [11]. While these data provide new opportunities for ACP discovery, several challenges persist, including the lack of automated tools for direct ACP prediction from raw meta-omics data, difficulties in identifying small open reading frames (smORFs) [12], and inconsistencies among existing prediction algorithms [9]. Additionally, current ACP prediction tools require peptide sequences as input [6,7], creating a computational bottleneck for microbial genome and metagenome samples. Together, these constraints hinder efficient and reliable ACP prediction, necessitating improved approaches.

To address these challenges, we developed MetaPepticon, a modular pipeline for large-scale ACP prediction from diverse inputs, ranging from raw sequencing reads to assembled contigs and peptide sequences. By integrating multiple ACP prediction tools within an automated workflow, MetaPepticon enables a consensus-based strategy to improve the reliability of candidate identification and generates a table of high-confidence ACPs for downstream analysis and is open-source and freely available at [13].

## Methods

### Pipeline overview

MetaPepticon is a modular and reproducible workflow for the discovery of ACPs from six input types: (i) single-organism shotgun sequencing (SG), (ii) shotgun metagenomics (MG), (iii) single-organism transcriptomics (ST), (iv) metatranscriptomics (MT), (v) assembled contigs (CO), and (vi) peptide sequences (PE). Implemented in Snakemake [14], each module runs within an isolated conda environment to ensure reproducible dependency management. Depending on the input type, the pipeline selectively activates modules for preprocessing, assembly, smORF prediction, ACP classification, toxicity filtering, and reporting (**Figure 1**).

**Figure 1.**
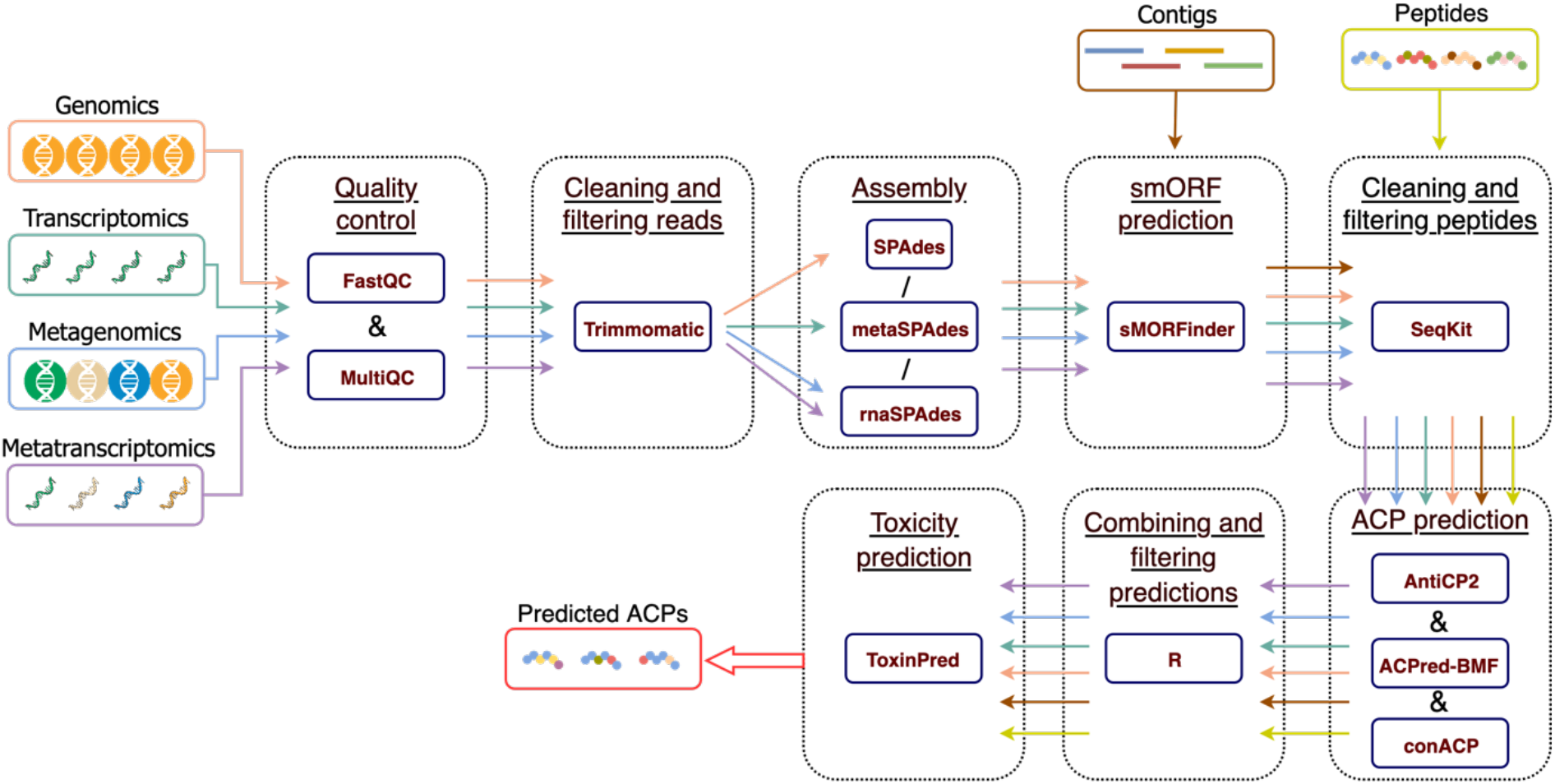
Overview of the MetaPepticon. MetaPepticon takes SG, MG, ST, MT, CO, and PE data as input and produces ACP and toxicity prediction results. The workflow consists of six modules, each optimized for specific input types. Key steps include: quality control, cleaning, and filtering (SG, MG, ST, MT); *de novo* assembly (SG, MG, ST, MT); smORF prediction (SG, MG, ST, MT, CO); peptide cleaning and filtering (SG, MG, ST, MT, CO, PE); ACP prediction (SG, MG, ST, MT, CO, PE); integration and filtering of ACP predictions (SG, MG, ST, MT, CO, PE); and toxicity prediction (SG, MG, ST, MT, CO, PE). The final output file contains peptide sequences alongside prediction results from each tool, customized by user-defined parameters.

### Input handling and configuration

Users initiate an analysis by placing input files into predefined folders (SG, MG, ST, MT, CO, PE) within MetaPepticon. Then, command line interface or graphical user interface can be used to generate a configuration file. Input file types and relevant parameters are automatically detected and presented to user. The configuration file exposes adjustable parameters including quality thresholds, assembly options, peptide length boundaries and consensus rules for ACP classification.

### Preprocessing and assembly

For raw sequencing reads, quality control is performed with FastQC [15], followed by adapter and quality trimming using Trimmomatic [16] (default: minimum Phred score = 25, minimum read length = 25 bp). Surviving reads are assembled with SPAdes [17] (for SG), metaSPAdes (for MG), or rnaSPAdes [18] (for ST and MT). A minimum contig length (default: 1000 bp) is applied to exclude short or low-confidence assemblies.

### smORF prediction and peptide filtering

smORFs are predicted using smORFinder [19], and the resulting peptides are filtered by length (default: 10–50 amino acids) with user-adjustable thresholds. Identical sequences are collapsed to a single representative to remove redundancy and then N-terminal methionine cleavage is performed.

### ACP prediction and consensus strategy

Candidate peptides are evaluated using three independent classifiers: AntiCP2.0 [20], ACPred-BMF [21], and ConACP [22]. MetaPepticon then applies a configurable consensus strategy, allowing users to retain ACPs predicted by at least one, two, or all three algorithms, depending on the desired stringency level.

### Toxicity filtering

All predicted ACPs are screened for potential toxicity with ToxinPred3 [23]. Sequences exceeding a configurable toxicity score (default: 0.38) are removed from the final dataset.

### Output and reporting

The pipeline generates a final table of predicted ACPs, including classifications from each prediction tool and toxicity assessments, as well as intermediate files, detailed logs, and quality control reports.

## Analyses

### Performance and characteristics of MetaPepticon

We evaluated the performance of MetaPepticon using experimentally validated ACPs from the CancerPPD2 database [24], together with a curated negative dataset of UniProt peptides excluding sequences annotated as antimicrobial or anticancer. After filtering for peptide length (10–50 amino acids), natural amino acid composition and redundancy, 1,031 peptides remained from CancerPPD2 and 1,767 from non-ACP dataset.

The heatmap of performance metrics revealed a trade-off between sensitivity and confidence (**Figure 2a**). The individual predictors differed in their strengths: Both AntiCP2 (precision 0.83, recall 0.73) and ConACP (precision 0.72, recall 0.81) achieved a balanced performance while ACPred favored sensitivity over confidence (precision 0.52, recall 0.91). At the least stringent consensus threshold (≥1 predictor), MetaPepticon maintained a high recall (0.96) while tolerating moderate precision (0.51, F1 0.67), capturing most true ACPs at the expense of some false positives. Increasing the threshold to ≥2 predictors improved precision (0.73) and specificity (0.82) while slightly reducing sensitivity (0.85), reflecting a shift toward higher-confidence candidate selection. The strictest consensus (3/3 predictors) maximized precision (0.91) and specificity (0.96) but substantially reduced sensitivity (0.64), illustrating the trade-off between confidence and coverage. Overall, the adjustable consensus thresholds enable flexible tuning of MetaPepticon, supporting broad discovery at lower thresholds (≥ 1) and high-confidence candidate selection at higher thresholds (≥ 2 or 3/3), depending on study objectives.

**Figure 2.**
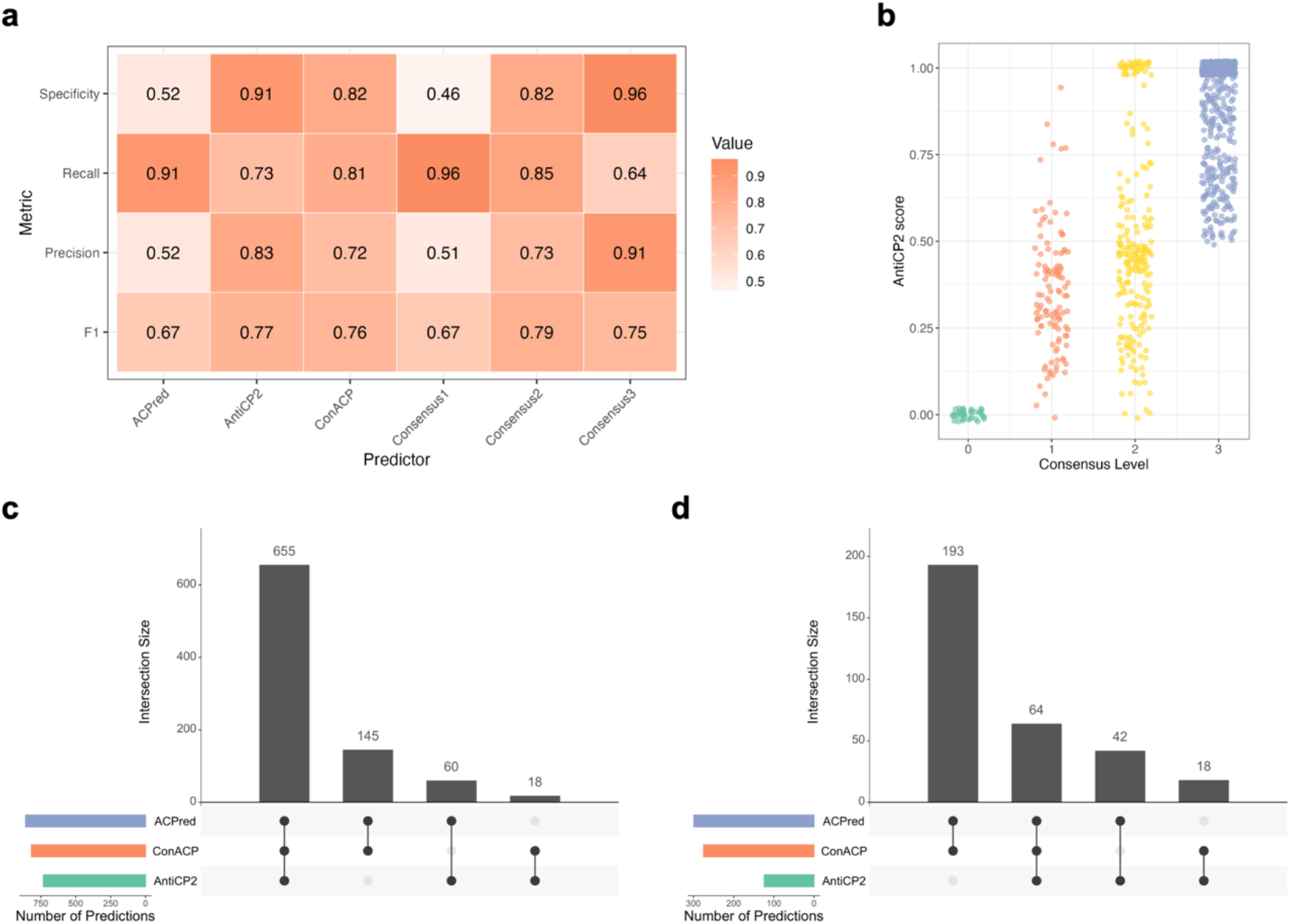
Performance and characteristics of MetaPepticon predictions. **(a)** Benchmark metrics for individual predictors (AntiCP2, ConACP, ACPred) and consensus thresholds (≥1, ≥2, 3/3) shown as grouped bar plots, displaying F1, precision, recall, and specificity values. **(b)** AntiCP2 prediction scores versus the number of predictors in agreement for validated positive peptides. **(c)** The overlap of predicted ACPs among AntiCP2, ConACP, and ACPred for positive peptides dataset. **(d)** The overlap of predicted ACPs among AntiCP2, ConACP, and ACPred for negative peptides dataset. Vertical bars indicate the number of peptides in each intersection, and horizontal bars show the total peptides predicted by each classifier.

AntiCP2 scores increased with higher consensus levels, with peptides predicted by multiple algorithms receiving stronger scores (**Figure 2b**). Moreover, AntiCP2 scores were positively correlated with the consensus of the other two predictors (Spearman’s ρ = 0.45, p = 2.2 × 10^−16^), supporting the use of a consensus-based approach. Analysis of prediction overlaps revealed that positive peptides were often identified by multiple classifiers (**Figure 2c**), while negative peptides showed little intersection (**Figure 2d**). Together, these observations indicate that increasing the consensus threshold reduces false positives among negative peptides, thereby enhancing the predictive specificity of the integrated pipeline.

### Prediction of candidate anticancer peptides from metagenomic datasets

To assess the performance of MetaPepticon, we analyzed ten gut metagenome samples from a previously published colorectal cancer cohort (BioProject PRJNA397219, Hale et al., 2018 [25]). For comparison, we processed the same samples with Macrel [26], an antimicrobial peptide (AMP) prediction tool that directly accepts raw metagenomic data (**Figure 3a**). Although Macrel is designed for AMP rather than anticancer peptide (ACP) prediction, it provides a relevant reference point for evaluating MetaPepticon’s end-to-end capability from raw metagenomic inputs. Because existing ACP predictors accept only peptide sequences, direct comparison at the metagenomic level was otherwise infeasible.

**Figure 3.**
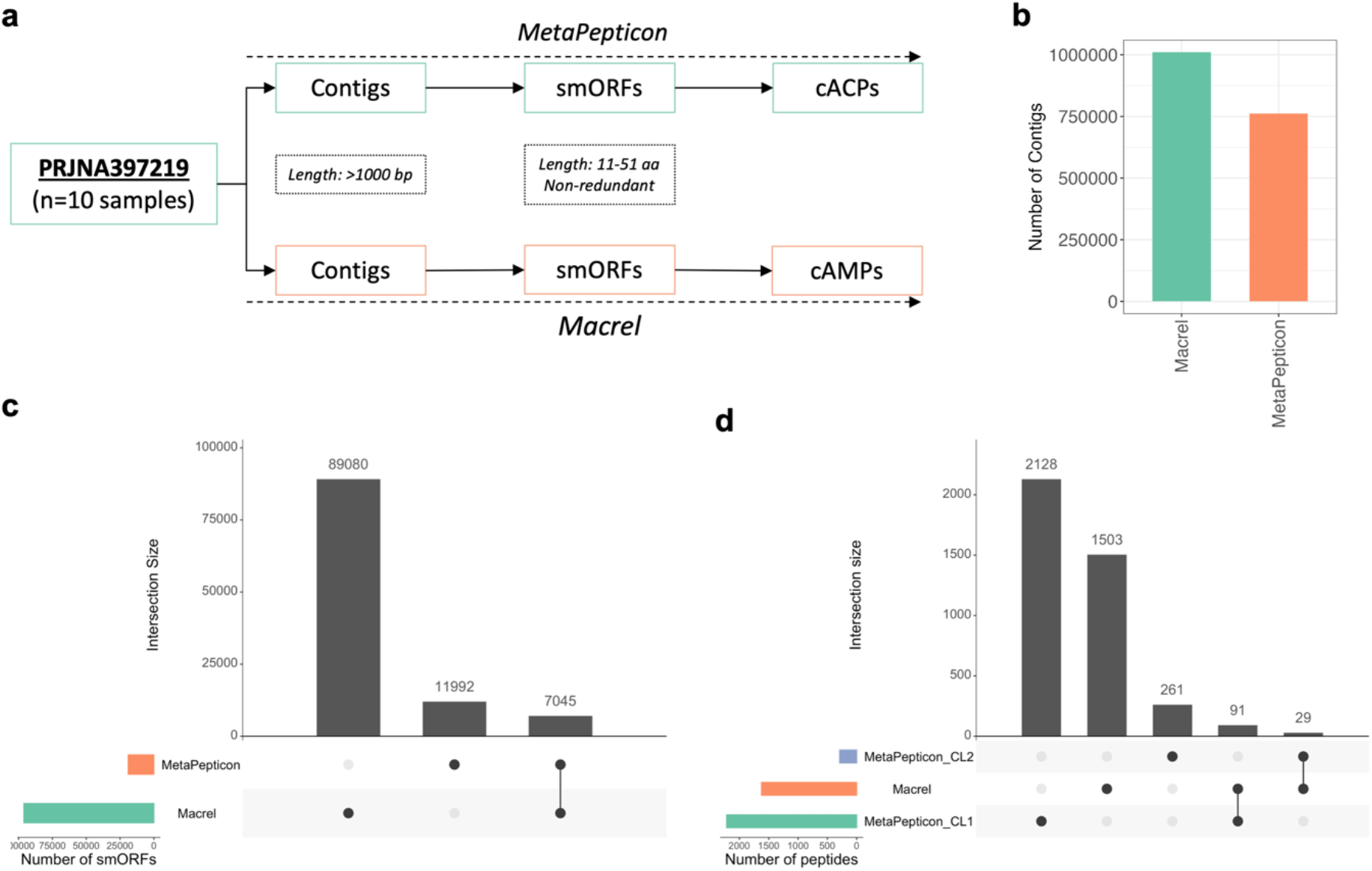
Comparative analysis between MetaPepticon and Macrel. **(a)** Ten gut metagenome samples were processed with both pipelines, each including de novo assembly, smORF prediction, and functional peptide classification (AMPs for Macrel, ACPs for MetaPepticon). **(b)** Total number of contigs generated by each tool. **(c)** Total and intersecting smORFs predicted by Macrel and MetaPepticon across the dataset **(d)** Total number of AMPs predicted by Macrel and ACPs predicted by MetaPepticon (at ≥1 predictor and ≥2 predictor consensus levels). Vertical bars indicate the number of smORFs/peptides in each intersection, and horizontal bars show the total number of predictions by each tool.

MetaPepticon generated 761,673 contigs using metaSPAdes, whereas Macrel produced 1,009,575 contigs (**Figure 3b**) using MEGAHIT [27]. At the smORF detection stage, MetaPepticon identified 11,992 unique smORFs, while Macrel predicted 89,080 (**Figure 3b**). This order-of-magnitude difference primarily reflects variation in ORF-calling strategies between two pipelines. Both rely on modified versions of Prodigal; however, Macrel employs a more permissive ORF-calling approach optimized for antimicrobial peptide discovery, whereas MetaPepticon uses smORFinder, which applies stricter length thresholds, ribosome-binding site modeling, and sequence-context filtering.

Among the smORFs detected by MetaPepticon, 7,045 (59%) were also identified by Macrel (**Figure 3c**). To assess whether this limited overlap was primarily driven by assembler choice, we re-ran Macrel using the metaSPAdes contigs generated within the MetaPepticon pipeline. Under this configuration, Macrel predicted 177,257 smORFs, of which 11,336 (95% of MetaPepticon’s smORFs) overlapped with the MetaPepticon results. These results indicate that assembler choice strongly affects the comparability of smORF sets and confirm that MetaPepticon recovers a conservative subset of high confidence smORFs.

At the functional peptide prediction level, Macrel reported 1,503 candidate AMPs (cAMPs), whereas MetaPepticon predicted 2,128 cACPs by at least one classifier. When a stricter consensus of at least two classifiers was required, the ACP set decreased to 261 (**Figure 3d**). Cross-comparison of functional predictions revealed limited overlap, with 91 peptides shared between Macrel AMPs and MetaPepticon ACPs at the single-predictor level and 29 remaining at the two-predictor consensus level (**Figure 3d**). This modest intersection suggests that antimicrobial and anticancer activity predictions, while based on partly overlapping physicochemical and sequence determinants, differ in the relative weighting and combination of these features.

Overall, this comparison underscores the principal sources of variability among existing tools, as differences in pipeline architecture markedly influence the search space. By incorporating multiple classifiers within an automated workflow, MetaPepticon offers a systematic approach that mitigates false positives through the use of a conservative smORF catalog and allows direct identification of cACPs from raw metagenomic data.

### Mining public microbial genomes and metagenomes for candidate anticancer peptides

We applied MetaPepticon to explore cACPs in large-scale publicly available microbial resources. Specifically, we analyzed 41,171 representative microbial genomes from the proGenomes3 [28] and 4,072,884 peptide sequences from the DBsmORF database [19]. Genomes were processed using the contig module and peptides with the peptide module of MetaPepticon under default parameters. From the proGenomes data, we detected cACPs by at least one of the integrated prediction algorithms while for DBsmORF peptides, 74,170 were predicted as cACPs (**Figure 4a**). Notably, 8,642 cACPs overlapped between the two datasets, consistent with the inclusion of RefSeq-derived genomes and Human Microbiome Project metagenomes in DBsmORF.

**Figure 4.**
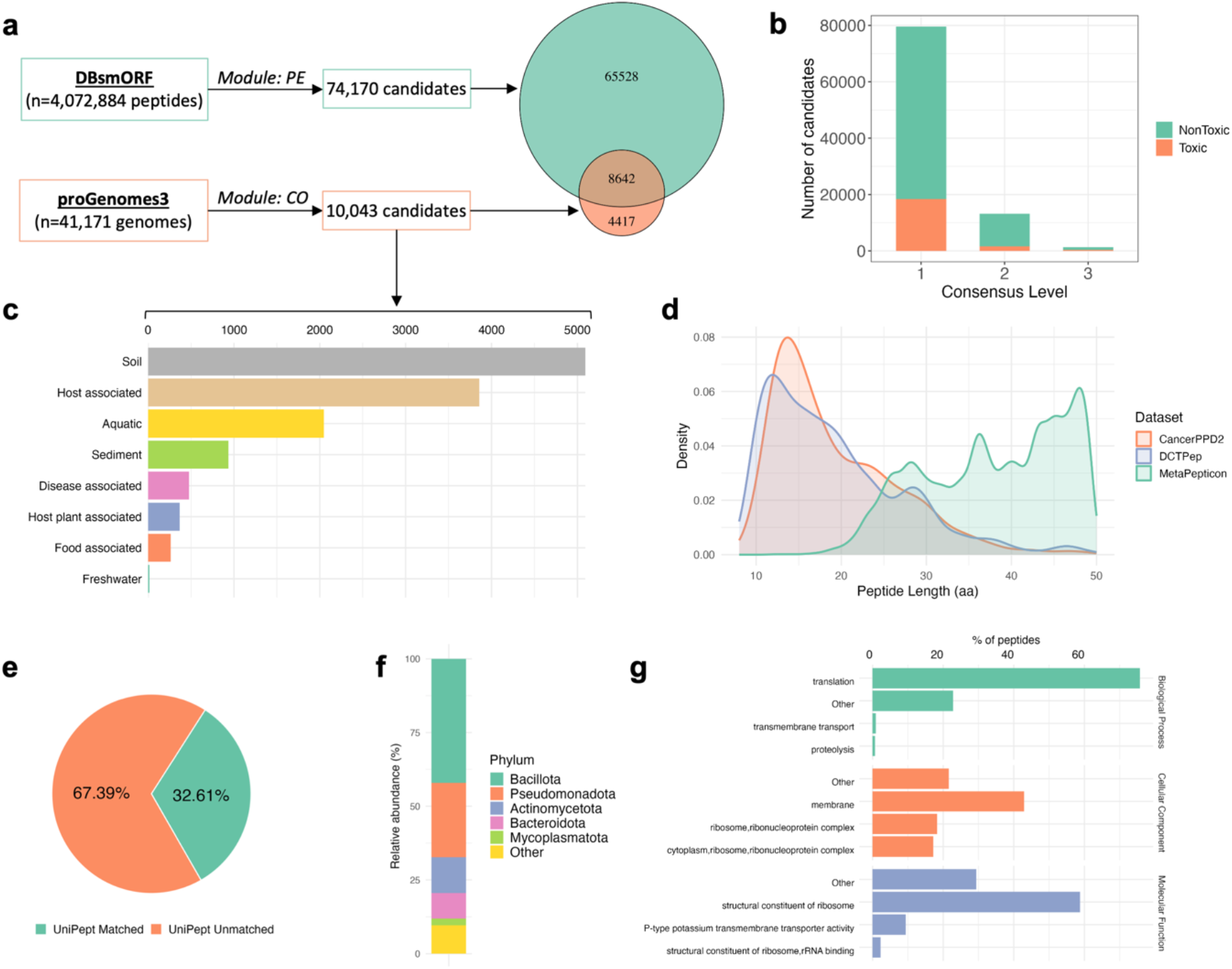
Discovery and characterization of novel cACPs using MetaPepticon. **(a)** Overview of the MetaPepticon pipeline applied to peptides from DBsmORF and contigs from proGenomes3. Venn diagram shows the number of cACPs identified from proGenomes3 versus DBsmORF, with intersection counts indicating shared cACPs. **(b)** The number of cACPs retained at different prediction stringency levels: ≥1 predictor, ≥2 predictors, and 3/3 predictors. Colors correspond to toxicity predictions; y-axis represents candidate counts on a linear scale. **(c)** The absolute number of candidates per habitat for proGenomes3-derived peptides. **(d)** Comparison of cACP lengths predicted by MetaPepticon with peptides from DCTPep and CancerPPD2 databases. **(e)** Venn diagram depicting the fraction of cACPs that could be annotated by UniPept at the ≥1 predictor consensus level. **(f)** Taxonomic distribution of annotated cACPs across bacterial phyla. **(g)** Gene Ontology (GO) assignments for annotated cACPs, shown separately for molecular function, cellular component, and biological process categories.

After merging results across both databases, we obtained 79,587 non-redundant novel cACPs. Comparisons with current ACP databases revealed no match for detected cACPs. Toxicity screening indicated that 61,145 of these were non-toxic. Applying more stringent consensus criteria reduced this number: requiring agreement by at least two prediction algorithms yielded 13,149 high-confidence candidates (11,610 non-toxic), while requiring consensus across all three algorithms identified 1,340 high-confidence ACPs (981 non-toxic) (**Figure 4b**).

Habitat-level analysis of ACPs predicted from proGenomes revealed that soil microbiomes contribute the largest absolute number of candidates followed by host-associated microbiomes (**Figure 4c**). The length distribution of cACPs peaked at 40–50 amino acids, differing from distributions observed in CancerPPD2 and DCTPep [29] databases (**Figure 4d**). This divergence likely reflects the novelty of microbial peptides, as peptides derived from metagenomes remain underrepresented in current ACP resources, a trend previously noted for cAMPs identified from global microbiomes [30].

To gain biological insight, we performed taxonomic and functional annotation of the high-confidence cACPs using UniPept [31]. Only 34% of peptides could be taxonomically annotated (**Figure 4e**), revealing diverse microbial origins dominated by *Firmicutes* (29.5%), *Actinobacteria* (23.3%), *Proteobacteria* (21.6%), and *Bacteroidetes* (13.1%) (**Figure 4f**). Functional profiling indicated broad molecular diversity, with ribosomal, translational, and membrane-associated peptides among the most enriched categories (**Figure 4g**). These results suggest that interactions with ribosomes and membranes may represent common mechanisms of action for microbially derived ACPs.

## Discussion

MetaPepticon is a modular bioinformatics pipeline designed for the discovery of cACPs from raw sequencing datasets, as well as contigs and peptide sequences. It streamlines the identification process by integrating genome assembly, open reading frame prediction, peptide annotation, and toxicity assessment within a single automated workflow. By incorporating multiple prediction algorithms, MetaPepticon allows users to balance broad discovery with high-confidence candidate selection, while toxicity predictions facilitate prioritization for experimental validation.

A key strength of MetaPepticon lies in its ability to bridge microbiome datasets with functional peptide discovery. The pipeline enables systematic mining of large microbial datasets, uncovering substantial numbers of novel cACPs across diverse microbial sources. Application to extensive public datasets demonstrates its capacity to identify previously uncharacterized peptides, expanding the repertoire of potential therapeutic candidates.

Although prediction accuracy depends on input quality, the consensus-based approach across multiple algorithms enhances reliability, providing a robust foundation for experimental follow-up. Future enhancements could include the integration of advanced machine learning models, structural modeling, and improved accessibility for users with limited bioinformatics experience, further refining functional predictions and broadening adoption.

Overall, MetaPepticon offers a reproducible and scalable framework for large-scale ACP discovery. By bridging raw sequencing data with peptide prediction, it supports systematic exploration of microbial peptide diversity and the identification of high-confidence candidates with potential therapeutic applications.

## Availability of source code and requirements

**Project name:** MetaPepticon

**Project home page:** https://github.com/arikanlab/MetaPepticon

**Operating system(s):** GNU/Linux

**Programming language:** Python, R and Shell

**License:** MIT

**Restrictions to use by non-academics:** No

## Acknowledgement

This work was supported by the Scientific Research Projects Coordination Unit of Istanbul University (Project Number: 41009).

## Authors’ roles

MA conceived the idea and designed the study. MA and AAE implemented the pipeline. MA performed analyses. MA and AAE wrote the manuscript. NT, GD, NÖŞ and MÇ provided critical feedback, reviewed and edited the manuscript. All authors approved the final version of the manuscript.

## Data availability

The public microbial genomes and peptide sequences analyzed in this study are available at https://progenomes.embl.de/download.cgi and http://104.154.134.205:3838/DBsmORF/, respectively. The gut metagenome samples analyzed are accessible under BioProject accession PRJNA397219.

## Competing interests

The authors declare no competing interests.

## Notes

**Relevant conflict of interest/financial disclosures:** Authors have no conflicts of interest to report.

**Funding:** This work was supported by the Scientific Research Projects Coordination Unit of Istanbul University (Project Number: 41009).

### Competing Interest Statement

The authors have declared no competing interest.

